# Three-dimensional study of spur morphogenesis in the flower of *Staphisagria picta* (Ranunculaceae) – from cellular level to organ scale

**DOI:** 10.1101/2023.07.20.549858

**Authors:** Pauline Delpeuch, Sophie Nadot, Katia Belcram, Antoine Plumerault, Catherine Damerval, Florian Jabbour

## Abstract

Floral spurs are invaginations borne by perianth organs (petals and/or sepals) that have evolved repeatedly in various angiosperm clades. They typically store nectar and can limit the access of pollinators to this reward, resulting in pollination specialization that can lead to speciation in both pollinator and plant lineages.

Despite the ecological and evolutionary importance of nectar spurs, the cellular mechanisms involved during spur development have only been described in detail in a handful of species, primarily with respect to epidermal cells. These studies show that the mechanisms involved are taxon-specific.

Using confocal microscopy and automated 3D image analysis, we studied spur morphogenesis in *Staphisagria picta* (Ranunculaceae) and showed that the process is marked by an early phase of dominant cell proliferation, followed by a phase of anisotropic (directional) cell expansion.

The comparison with *Aquilegia*, another taxon of Ranunculaceae with spurred petals, revealed that the convergence in form between the spurs of both taxa is obtained by partially similar developmental processes. The analytical pipeline designed here is an efficient method to visualize in 3D each cell of a developing organ, paving the way for future comparative studies of organ morphogenesis in multicellular eukaryotes.

**Highlight:** A new method of 3D analysis of plant tissues at the cellular level revealed that spur morphogenesis in *Staphisagria picta* is marked by an early phase of dominant cell proliferation, followed by a phase of anisotropic cell expansion. Floral spur development is analysed for the first time quantitatively, taking into account all tissues composing the organ, namely epidermis and parenchyma.

## Introduction

The way in which similar forms or functions may be acquired independently through different or similar developmental processes, involving homologous genes or completely different genetic mechanisms is an important question in evolutionary biology. Floral nectar spurs are invaginations of various dimensions borne on perianth organs. They have evolved repeatedly in angiosperms, in various clades, on petals (for example in some lineages of orchids and Lamiales, in *Viola* (Violaceae, Malpighiales), or *Valeriana* (Caprifoliaceae, Dipsacales)) or sepals (for example in *Tropaeolum* (Tropaeolaceae, Dipsacales) (Ronse De Craene and Smets, 1995)) (Figure 1). They usually store nectar, and may restrict access to this reward, filtering the most efficient pollinators. Therefore their presence could be linked to specialization in pollination and possibly lead to speciation in both pollinator and plant lineages (Whittall and Hodges, 2007). Despite the ecological and evolutionary importance of floral nectar spurs, the cellular mechanisms taking place during spur development have been described into detail only in a few species, often considering only epidermal cells. Development classically consists of two phases (cell proliferation and cell expansion and differentiation which in plants are most generally segregated (Walcher-Chevillet and Kramer, 2016)), whose relative proportion and timing differ among clades. In *Aquilegia* (Ranunculaceae, Ranunculales), detailed observations of the spur epidermis suggested that the shape of the mature spur results from a combination of cell proliferation at early stages of development, followed by anisotropic cell expansion allowing spur elongation. The comparative study of spur development in four *Aquilegia* species revealed that anisotropic cell expansion accounts for the differences in spur size among species (Puzey *et al*., 2012). A similar study conducted in *Linaria* (Plantaginaceae, Lamiales) suggested that differences in spur length are better explained in this genus by differences in the number of cells resulting from the initial phase of cell proliferation (Cullen *et al*., 2018). The variation in the duration of the phase of cell division supports the hypothesis that changes in the activity of cell cycle genes and their regulators may be involved in the evolution of the nectar spur shape and dimension. Like for *Aquilegia*, *Valeriana rubra* (basionym of *Centranthus ruber,* Caprifoliaceae, Dipsacales), anisotropic cell expansion of epidermal cells plays a predominant role in spur development (Puzey *et al*., 2012; Mack and Davis, 2015).

**Figure 1:**
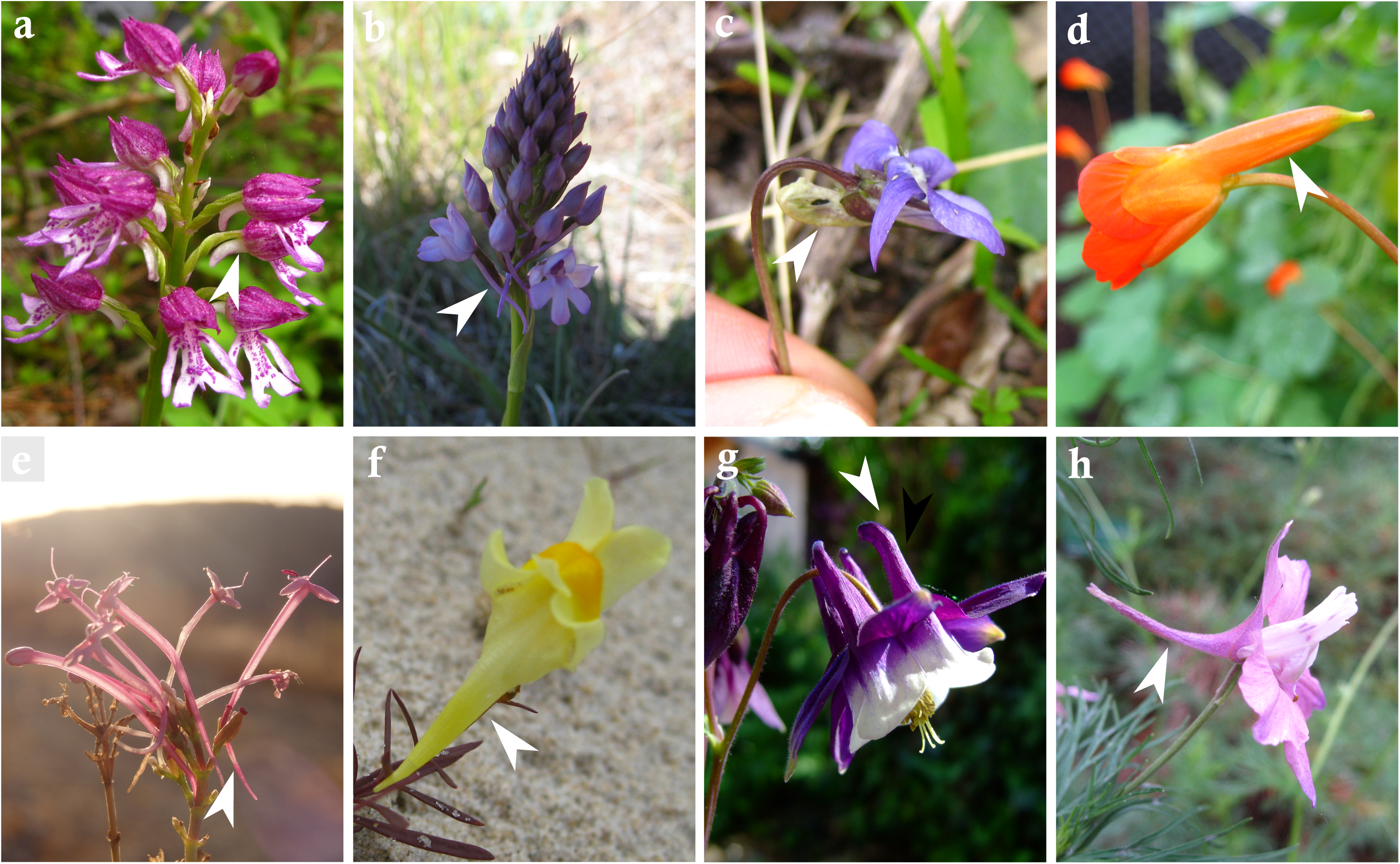
Spur diversity in angiosperms (a) *Orchis militaris*, (b) *Anacamptis pyramidalis*, (c) *Viola riviniana*, (d) *Tropaeolum tuberosum*, (e) *Valeriana erotica*, (f) *Linaria vulgaris*, (g) *Aquilegia vulgaris*, (h) *Delphinium ajacis*. Photographs: Sophie Nadot except (g): Wikimedia © Aiwok.

The Ranunculaceae family comprises approximately 55 genera - ca. 2,500 species - that display a great floral diversity, particularly at the perianth level, which may be composed of both sepals and petals, or only sepals (reviewed in (Carrive *et al*., 2020)). Spurs evolved repeatedly in Ranunculaceae: twice on petals (in the stem lineages of the genus *Aquilegia* and of the tribe Delphinieae) (Figure 2), and twice on sepals (in the stem lineages of the tribe Delphinieae and of the genus *Myosurus*) (Hiepko, 1965; Kosuge and Tamura, 1988; Kosuge, 1994; Hodges, 1997; Erbar *et al*., 1999; Endress and Matthews, 2006; Delpeuch *et al*., 2022). The diversity in spur shape observed in the genus *Aquilegia* makes this genus an ideal model to study how the interactions with pollinators may have shaped floral morphology (Whittall and Hodges, 2007), and also to identify the mechanisms of spur development and the genes possibly involved (Ballerini *et al*., 2019, 2020; Zhang *et al*., 2020b). Spur morphogenesis begins with the formation of a depression in the center of each petal primordium, that further expands to form a hollow and narrow invagination (Tucker and Hodges, 2005; Ren *et al*., 2011). At the cellular scale, the formation of the depression involves a short period of localized cell divisions that stop progressively from the petal tip to the the site of initiation, then the deepening of the cup is achieved by anisotropic cell expansion (Puzey *et al*., 2012). Cell proliferation is controlled by hormone signalling, principally involving auxine response genes (Yant *et al*., 2015; Zhang *et al*., 2020). In flowers of Delphinieae, the dorsalmost petals are spurred and nectariferous. These petals are nested within the spurred sepal (Jabbour and Renner, 2012b), and their morphology (Figure 2) and development have been extensively studied *(Jabbour et al., 2009*; Chartier *et al*., 2016; Chen *et al*., 2018; Jabbour *et al*., 2021). It is a case of synorganization, that is to say the intimate structural connection of two or more neighbouring floral organs forming a functional system (*i.e*., a hyperorgan) (Specht and Bartlett, 2009; Jabbour *et al*., 2021). The length of the inner spurs determines the range of pollinators able to collect nectar. The genus *Staphisagria* is sister to the remaining Delphinieae and includes two species, namely *S. picta* and *S. macrosperma* (Léotard, 2002; Jabbour and Renner, 2011). This genus was used to describe the development and structure of the nectariferous hyperorgan (Jabbour *et al*., 2021). A depression is initiated at the primordium stage, deepens, and eventually forms a curved spur (Zalko *et al*., 2021). However, the cellular mechanisms involved - in terms of localization and duration of cell proliferation and expansion - remain undescribed.

**Figure 2:**
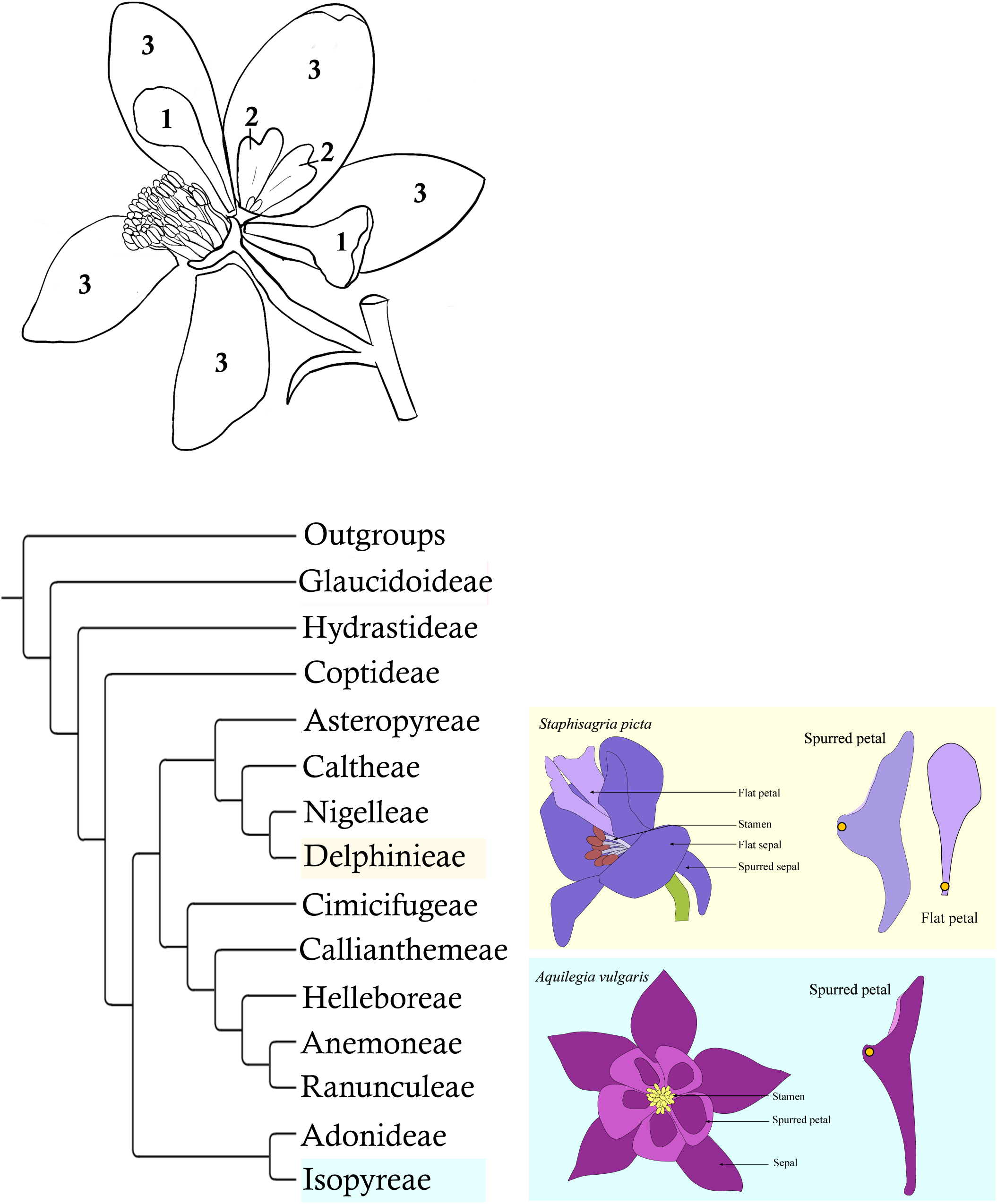
Floral morphology of *Staphisagria picta* and simplified phylogeny of Ranunculaceae, with tribes characterised by spurred flowers highlighted Ranunculaceae tribes. A: Structure of the flower of *Staphisagria picta*: (1) flat petals, (2) spurred petals, (3) sepals. B: Phylogenetic relationships among Ranunculaceae tribes (based on Zhai *et al.,* 2019); tribes including taxa characterized by flowers with spurred petals are highlighted in yellow (Delphinieae) and blue (Isopyreae). On the right hand of the figure: flowers of *Staphisagria picta* (top) and *Aquilegia vulgaris* (bottom) and details of their petals. The orange dot indicates the insertion point of the organ on the floral receptacle. The corolla of *S. picta* consists of two flat lateral petals and two spurred dorsal petals. The pair of dorsal petals is nested within the spurred sepal.

The aim of the present study is to finely explore spur morphogenesis by using the species *S. picta* as a model to address the following questions: is spur length and curvature mostly explained by cell proliferation or by cell expansion? Does the independent dual origin of spurred petals in Ranunculaceae result from a convergence in shape or also from a convergence in the pattern of cellular processes? Because of the phylogenetic affinity of *Staphisagria* with *Aquilegia* (both genera belong to the same family), we expect anisotropic cell expansion to account mostly for spur growth, cell division being restricted to the earliest developmental stages. To test this hypothesis, we relied on the MorphoLibJ library, available on the open-source platform for biological-image analysis Fiji (Schindelin *et al*., 2012; Legland *et al*., 2016), a collection of generic tools dedicated to the analysis of plant cells on 3D images. We characterized spur morphogenesis in *S. picta* using confocal microscopy and described the three-dimensional characteristics of cells (including epidermis and parenchyma) at successive developmental stages. The results are compared with those previously obtained in *Aquilegia* (Puzey *et al*., 2012)*, Centranthus* (Mack and Davis, 2015), and *Linaria* (Cullen *et al*., 2018), focused on epidermal cells.

## Materials and methods

### Plant material

Seeds were obtained from the French National Museum of Natural History (MNHN). Reference herbarium specimens corresponding to plants grown from the same set of seeds are kept at P herbarium (MNHN) (barcode P04023155, http://coldb.mnhn.fr/catalognumber/mnhn/p/p04023155, and P04023156, http://coldb.mnhn.fr/catalognumber/mnhn/p/p04023156). Floral buds were sampled from plants grown at the Jardin Botanique de Launay (Orsay, France) in September 2021, and fixed in FAA (90% alcohol 70%, 5% formaldehyde, 5% acetic acid). The corolla of *Staphisagria* species comprises four fully-developed petals. At anthesis, the two lateral petals are flat, whereas the two dorsalmost petals are spurred (Figure 2) and nested within the hollow sepal. The nectariferous tissue is located all along the inner epidermis of the spur (Zalko *et al*., 2021). Floral buds of *Staphisagria picta* were collected at 10 successive developmental stages, from stage 01 (bud sampled after the resumption of development, the petal is flat) to stage 10 (mature petal). The development of reproductive organs was used as a reference to calibrate the developmental sequence (Supplementary Fig.S1, adapted from (Delpeuch *et al*., 2022)). Buds representative of each of the ten developmental stages, covering the whole developmental sequence, were selected (one bud per stage), and one dorsal spurred petal – left or right – was dissected for further analysis (Figure 3).

**Figure 3:**
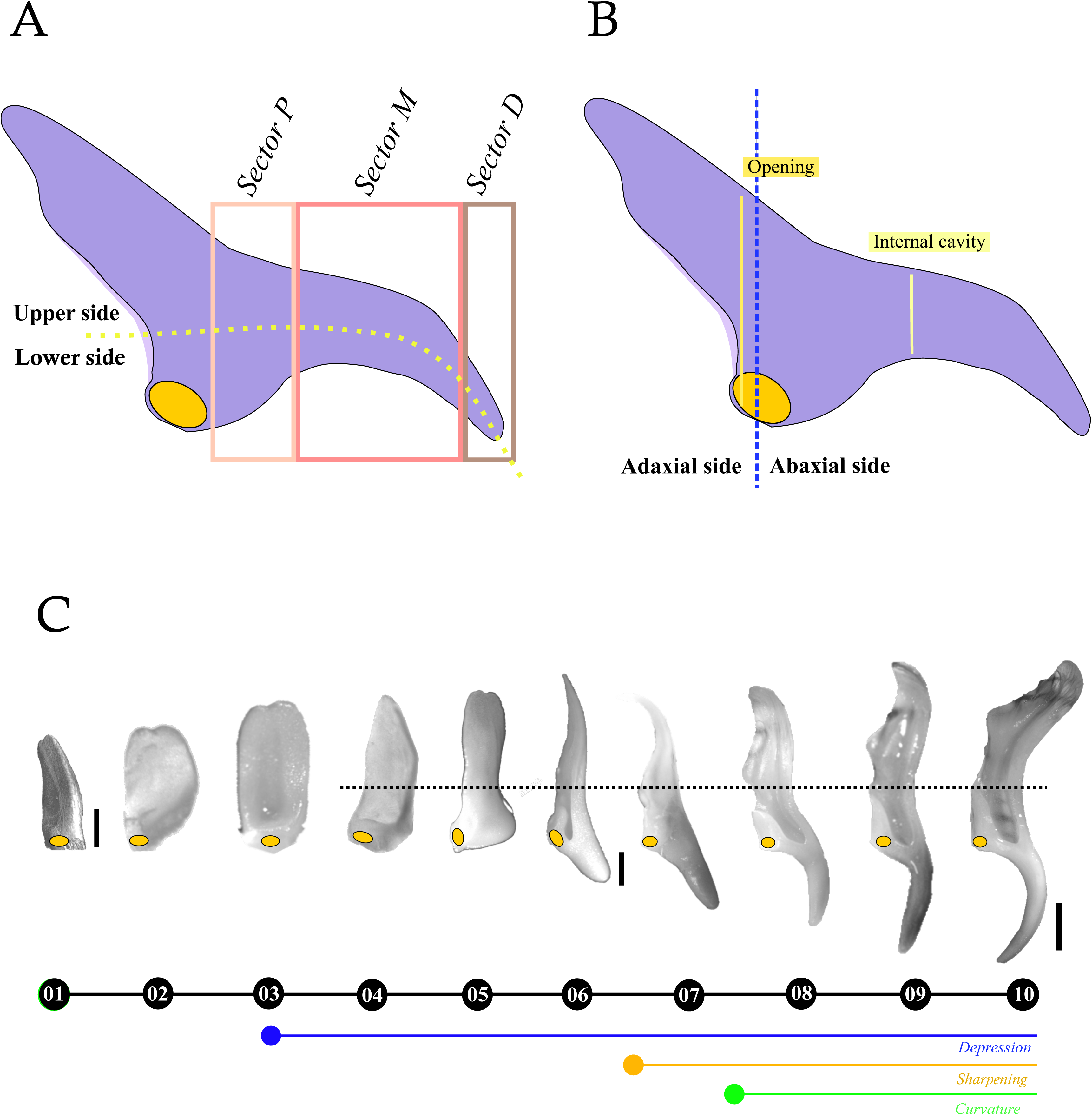
Orientation of petals in space and along the developmental sequence. (A) Definition of sectors: proximal, median, distal and lower and upper sides, (B) Definition of the spur opening and internal cavity, (C) photographs of petals along the developmental sequence. The orange dots indicate the petal insertion point on the floral receptacle. Scale: stages 01– 05: 150 µm, 06: 1mm, 07–10: 2mm).

### Confocal microscopy

Tissues were treated as described by Schaefer *et al*. (2017). Cell walls were stained with fluorescent brightener 28 as described by Belcram *et al*. (2022) with minor modifications as follows. Samples fixed in FAA were transferred in ethanol 70%, and then progressively rehydrated in 50% ethanol, followed by 10 minutes in 30% ethanol. They were incubated in 0.2 N sodium hydroxide/1% SDS for two hours at room temperature and rinsed in water. Samples were then simultaneously cleared and stained by an incubation overnight in a clearing solution (25% urea, 15% deoxycholate, 10% xylitol in distilled water) with the addition of 0.1% fluorescent brightener 28 (the stock solution is a 1% solution with one drop of sodium hydroxide 10 N to allow complete dissolution). Samples were rinsed in the clearing solution without the staining solution; and mounted in Citifluor AF1 (Agar Scientific). The image acquisitions were made with an inverted Zeiss Observer Z1 spectral confocal laser microscope LSM 710 and with a 25x objective (LD LCI Plan-Apochromat 25x/0.8 Imm Korr DIC M27). Fluorescence of the Fluorescent Brightener 28 dye was recorded using a 405-nm excitation and a selective emission window of 410–485 nm. Each sample was imaged as a Z-stack (of longitudinal sections) encompassing the entire thickness of the spur. We used a voxel size of 0.35 × 0.35 × 0.5 μm for the first five stages and 1.1070 × 1.1070 × 1.1 μm for the last five.

### Scanning electron microscopy

Buds of *Staphisagria picta* at mature stage (stage 10) and between stages 06 and 07 were fixed in FAA. They were placed in ethanol 70% the day before dissection. Petals were extracted and longitudinal and tangential sections were made under a stereoscope (Nikon SMZ 745T). The dissected petals were dehydrated in absolute alcohol and dried using an Emitech K850 critical-point dryer (Quorum Technologies), mounted on aluminum stubs with colloidal graphite, sputter-coated with platinum (60 s of metallization) using a JFC-1200 fine coater (JEOL), and observed using an SU3500 scanning electron microscope (Hitachi).

### Imaging strategy

Buds at early developmental stages (stages 01 to 05) were imaged as a whole, without isolating the developing petals, because of their very small size. In larger buds (stages 06 to 10), petals were extracted, and the spur was isolated and tagentially sectioned. Each side was imaged separately. The petals were oriented in space according to their insertion on the floral receptacle. For the largest samples (stages 06 to 10), cells were first analysed along the entire imaged spur. In a second step, the spur was divided into three sectors, corresponding respectively to the proximal zone (sector P: closest to the receptacle), the median zone (sector M), and the distal zone (sector D: tip), and the imaging and analyses were repeated for each sector. The sectors were defined along the corresponding longitudinal axis. They were first defined on the mature petal, and transferred to the other stages, adapting their size. Analyses were performed on the upper side and lower side of the spurs of stages 06 to 10 separately, to study the curvature of the spur in detail. The lower side was defined as the part of the spur located in the prolongation of the zone of insertion to the floral receptacle, and the upper side as the part opposite the zone of insertion to the floral receptacle (Figure 3).

### Segmentation processing

Images were subdivided into smaller parts to fit in the hardware memory using the software Fiji (Schindelin *et al*., 2012). First, we applied on the images the pre-processing of PlantSeg (Wolny *et al*., 2020), a pipeline for volumetric segmentation of plant tissues into cells. This pre-processing employs deep learning, in particular a convolutional neural network to predict cell boundaries. Second, all images were blurred and segmented with the Morpholib package (Legland *et al*., 2016), selecting parameters “catchment basins method” with a tolerance of 0.5. Parameters of the segmentation were chosen empirically by performing manual segmentations tests. Because of the size and number of acquired data, automation of the segmentation process launched with a homemade Python script was necessary.

### Visualization and analysis of segmented petals

A script written in Python language using the package “plotly” allowed us to visualize the result of the segmentations. All segments were reassembled using their coordinates and each cell was displayed according to its x, y, and z coordinates. Petals were reconstructed in three dimensions and the characteristics of cells (volume, sphericity, and orientation of aspheric cells) were visualised using different colours, and further analysed quantitatively. Cell density in each sector is then simply the inverse of the average cell volume. The volume of the organ or sector is estimated by the total volume of the cells, independently of the hollow part of the spur.

The data extracted from image analysis provide information on the alignment of cells along a given axis, for example the spur longitudinal axis. The value of the angle 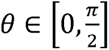 between cell orientation vector and the axis was used as a measure of the alignment of the cells with the axis. To interpret the result, the theoretical distribution of angles for an isotropic distribution of direction is shown in black on the figures. This distribution *p*(*θ*) can be calculated by considering that the surface element of the unit sphere spanning *θ* to *θ* + *dθ* is *ds* = *dθ* sin(*θ*)2π, thus *p*(*θ*) = *C* sin(*θ*), with *C* a constant to be determined. Because *p* is a probability distribution we have 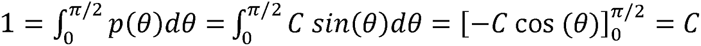, i.e. *C*= 1 and *p*(*θ*) = sin (*θ*).

### Post-processing

Pre-processing with PlantSeg amplifies the noise of the acquisition in the background. As a result, “false cells” may appear when performing the segmentation. To discard most of these “false cells”, the background was removed by applying a mask on each petal. Such mask was obtained by running segmentation using CLAHE instead of PlantSeg, which is less precise but has the advantage of being less prone to generating “false cells” in the background. To create the 3D mask, the volume was partitioned into smaller cubes, the number of cells in each cube was calculated, and cubes with a number of cells under a threshold empirically chosen were masked. This process allowed us to discard regions outside the petal without discarding regions of the petal itself. The mask was then applied on the predictions of the PlantSeg segmentation to remove “false cells”.

Cell outliers, i.e. the 5% largest and smallest cells in terms of volume, were filtered out. To visualize the interior of the petals, we relied on the opacity of the dots or on virtual sections.

The code that allowed the automation of the segmentations and the visualization of the data is available on github [https://github.com/paulinedlpch/morphogenesis].

## Results

### Stages of floral development

Flower development of *Staphisagria picta* begins with the initiation of the five sepal primordia, followed by petal initiation. Petal development is arrested shortly after stamen initiation. It resumes after the initiation of carpel primordia. In the present study, we examined only the developmental stages that take place after the developmental stasis of petals. At stages 01 and 02, the petal is flat, thick, and curved, slightly lobed in the distal part. A depression in the blade appears between stages 02 and 03, in the shape of a broad pocket. The depression deepens, developing into a spur between stages 05 and 07. Between stages 07 and 08, the spur becomes slightly curved in its proximal part (close to the receptacle). The rest of the spur becomes curved between stages 08 and 09. The spur lengthens throughout the whole development. At stage 10, the mature spur is curved, *ca.* 7.5 mm long and 2 mm wide at the opening.

### Cellular characteristics during petal development

On raw images obtained by confocal microscopy and 3D visualizations, epidermal cells appeared larger and had a more regular shape than inner cells (Figure 4AB). The inner epidermal layers are evenly organized. Spherical, small and disordered cells could be distinguished at stage 01 at the position of the future lobes. From stage 02, the central zone presents evenly ordered and slightly elongated cells. Large and irregular cells were observed towards the insertion point of the petal. From stage 03, three rows of elongated cells oriented along the longitudinal spur axis were observed, which could correspond to the vasculature (Figure 4CD). These rows of elongated cells multiply, branching throughout the spur as development proceeds.

**Figure 4:**
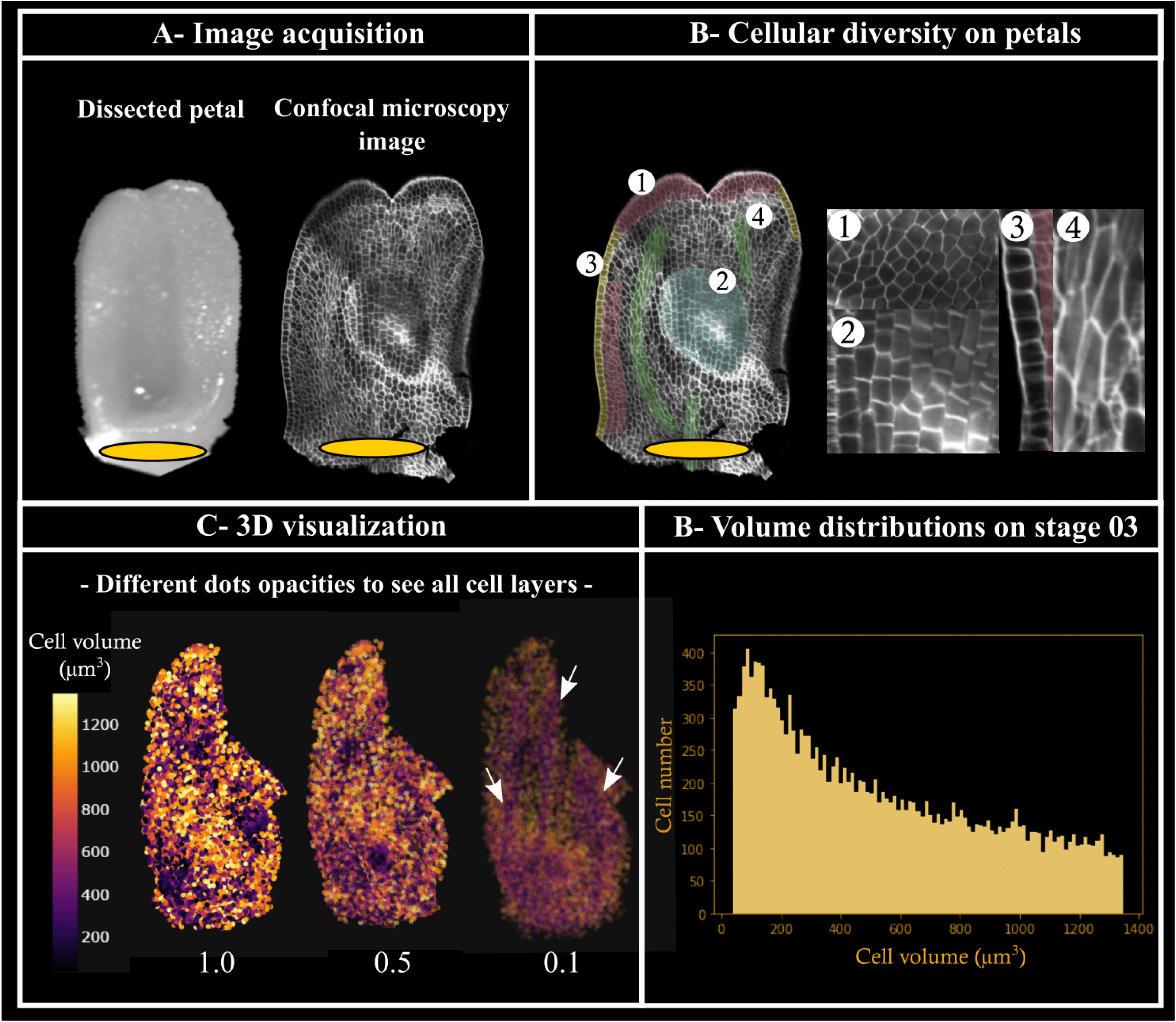
Different steps in the visualization of results illustrated by adaxial views of stage 03 petals. A: confocal microscopy images, only one cell layer is shown but it is possible to move layer by layer within the petal. B: different cell shapes: (1) small parenchyma cells, (2) large parenchyma cells, (3) regular, large epidermal cells present in a single cell row, (4) elongated cells belonging to the vascular system. C: visualization of the results by reconstruction of the petals in 3D with *in silico* staining of the cells according to their volume. The three lines corresponding to the vascular system are indicated by white arrows. D: histogram of cell volume distribution. The orange ovals indicate the petal insertion point on the floral receptacle.

### Average cell volume in whole spurs

Based on the variation of the average cell volume during spur development, two phases can be defined. From stage 01 to 06 (i.e. during spur formation), mean cell volume remains relatively stable (on average between 550 and 900 µm^3^) while cell number and total volume of the spur increase, suggesting cell proliferation as the main process. From stage 07 to 10 (i.e. during spur growth), the mean cell volume increases continuously (until on average 5,000 µm^3^ (stage 07) to 20,000 µm^3^ (stage 10), in parallel with the total spur volume (Figure 5A). This observation is consistent with cell expansion as the main cellular process at work during this second phase. In other words, between the two phases there is a 36-fold increase in volume or a 3.3-fold increase in size. Note that the stage 09 seems particular, in which the cell volume does not follow the overall trend.

**Figure 5:**
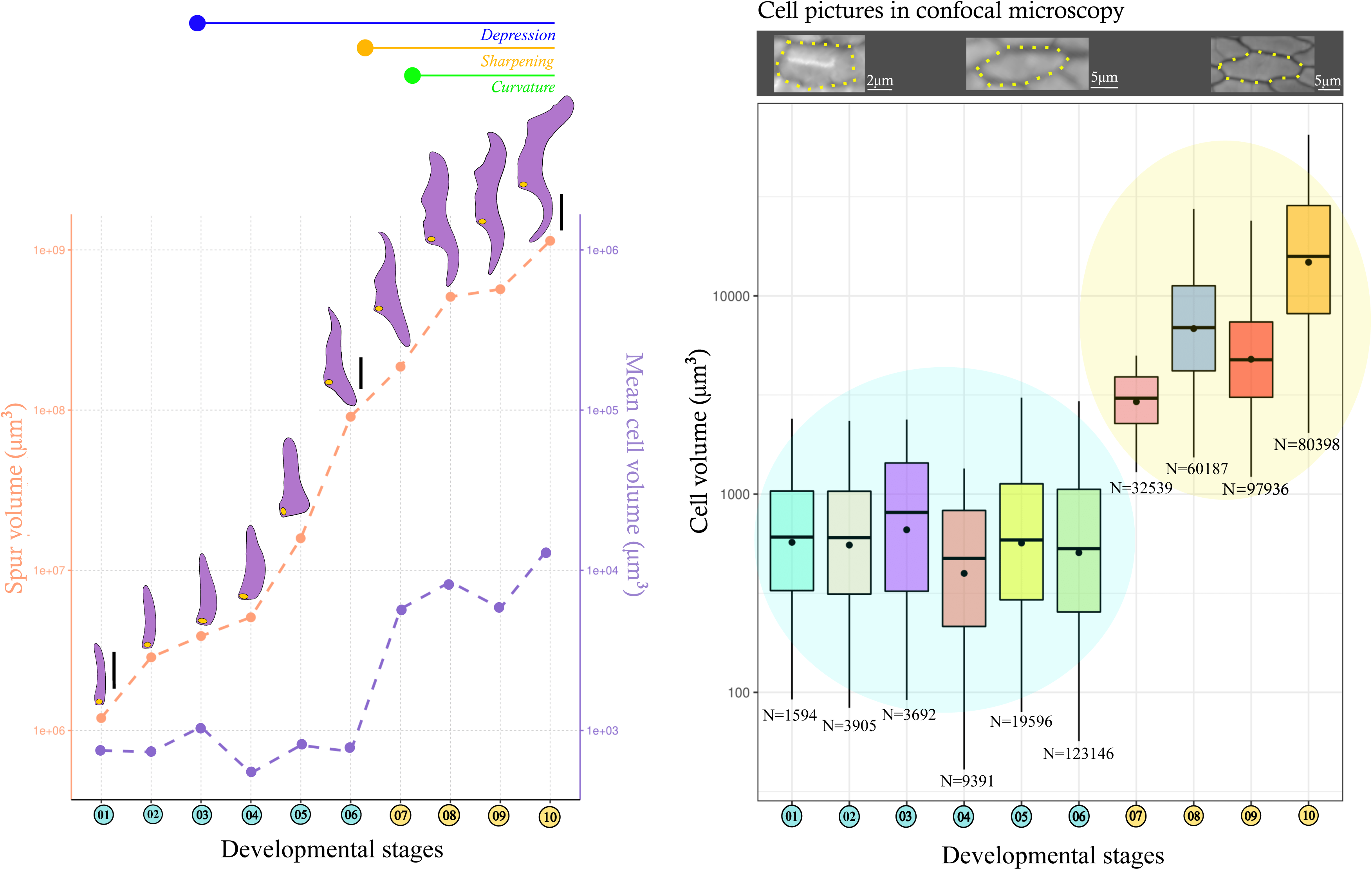
Variation of spur and cell volume during development. Left, description of the petal shape, and evolution of mean cell volume compared to whole spur volume during development. All values are in logarithmic scale. Scale: stages 01–05: 300 µm, 06: 2 mm, 07–10: 4 mm). The yellow dots represent the area of insertion to the floral receptacle. Right, boxplots represent the volume of cells at each stage. Blue and yellow areas represent the first and second phase of development, respectively. N: cell number per stage.

At stage 06, the spur has six times more cells than in the previous stage (stage 06: 123,146 cells versus stage 05: 19,596 cells). At the following stage, there are 3 times less cells (Stage 07: 32,539 cells) (Figure 5B, Supplementary Fig. S2).

### Average cell volume in sectorised spurs

Overall during development, an increase in cell volume is observed in the spur. However, at each developmental stage, cell volume differs significantly among proximal, median and distal sectors. The mean cell volume decreases and cell density increases towards the tip of the spur. Cell density of the distal cells (sector D) is the highest and the cells are the smallest (Stage 10 - Figure 6).

**Figure 6:**
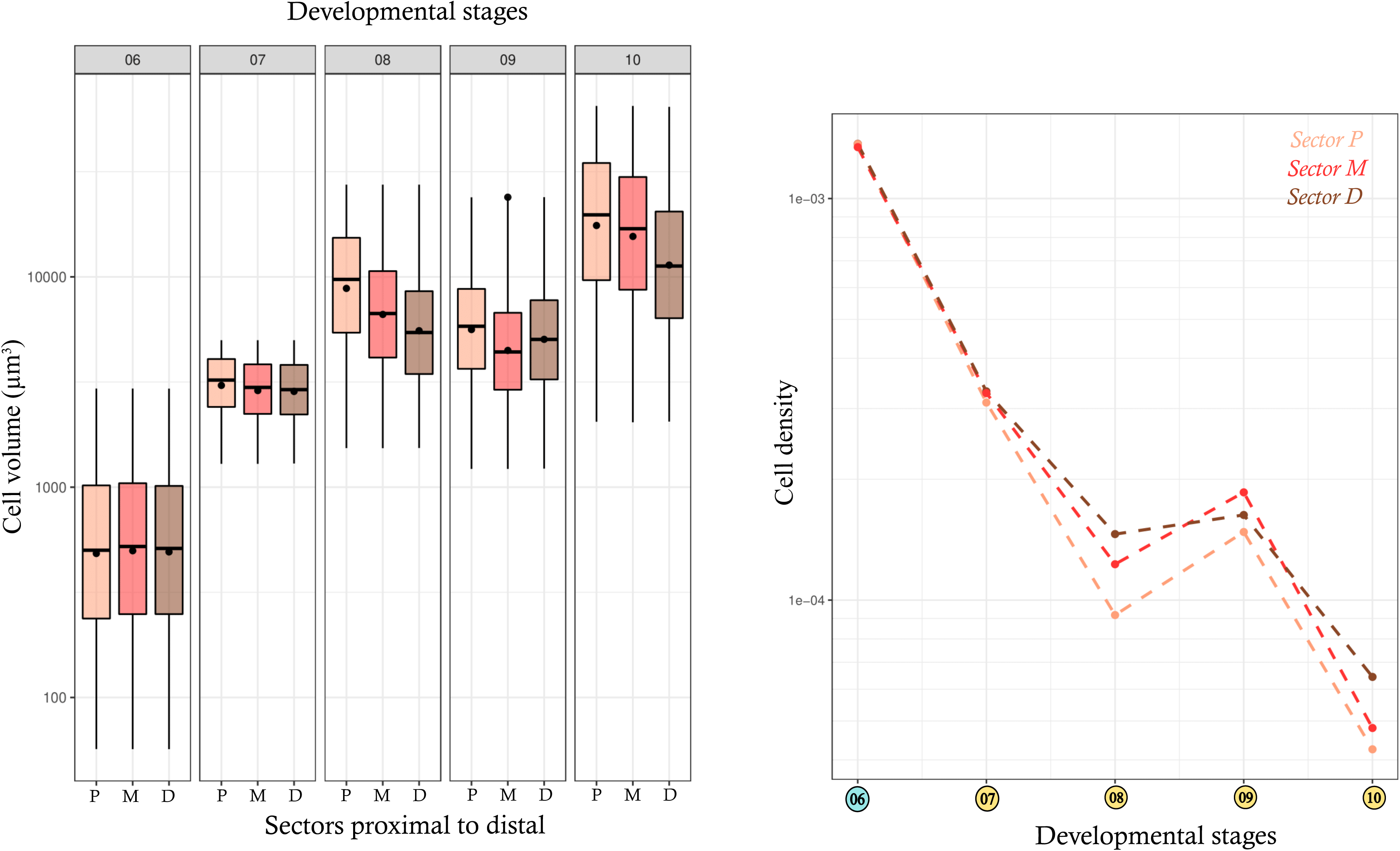
Variation of cell volume and density during spur development. Left, boxplots representing cell volume (µm^3^) according to stages and sectors. Right, representation of cell density on the proximal (P), median (M) and distal sector (D) along the developmental sequence. All values are in logarithmic scale.

### Cellular anisotropy

In the spur, cell orientation varies throughout development (Figure 7). They tend to be orthogonal to the longitudinal axis at stages 01 and 10. At stages 02, 04, 05, and 06, cell orientation follows the direction of the longitudinal axis. At stage 03 and stages 07 to 09, cells are distributed in two intermingled main populations depending on their orientation. The orientation of one cell population follows the longitudinal axis, whereas the orientation of the other population is orthogonal to the longitudinal axis. These two populations are visible on SEM images of tangential and longitudinal sections of spurs on mature petals or petals collected at stages 07 (Supplementary Fig. S3).

**Figure 7:**
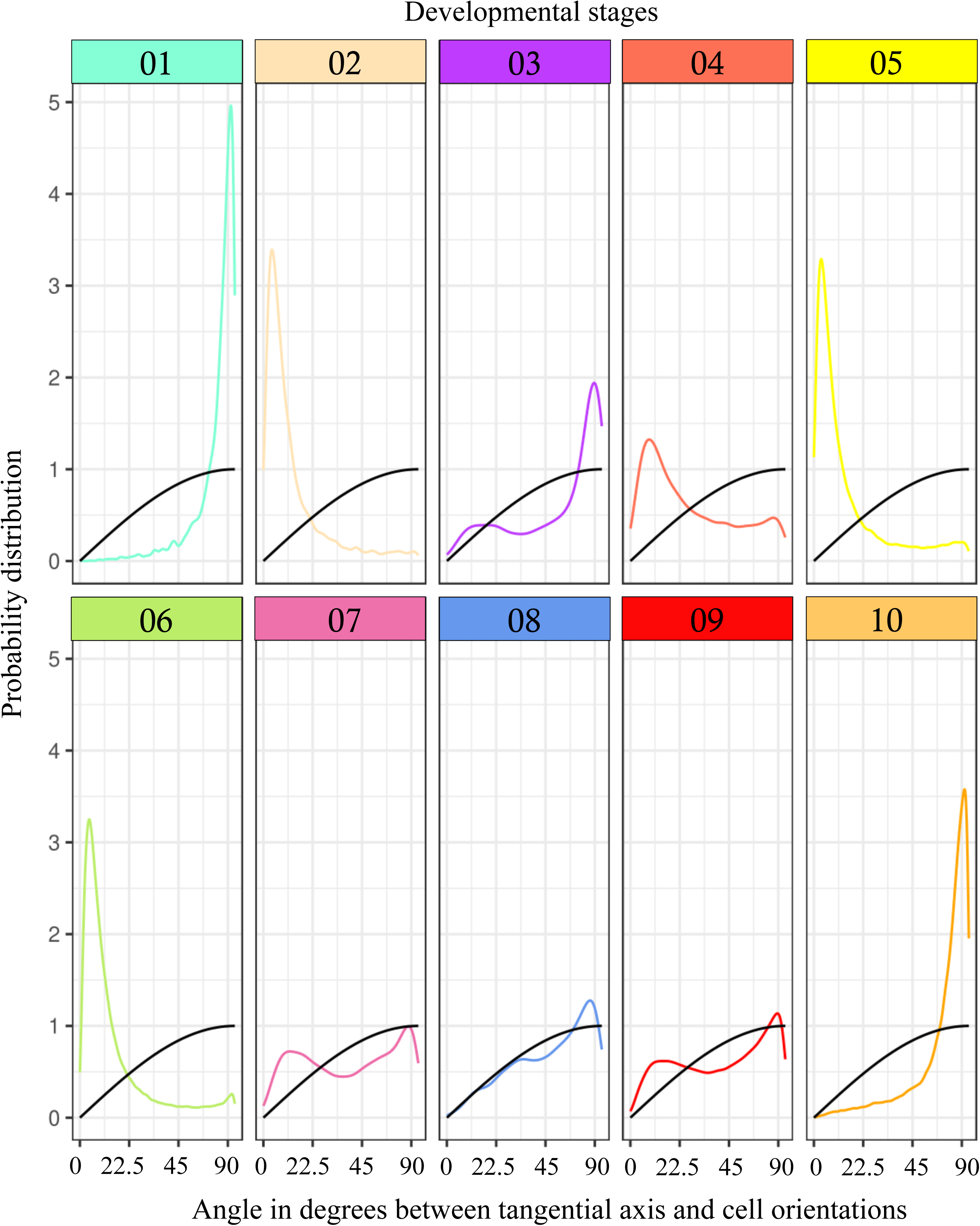
Quantification of cellular anisotropy during spur development. Plots representing the deviation angle of the directional axis of cells from the direction of the longitudinal axis of the petal for the spur. The black curve represents the distribution of randomly spheric oriented cells. Significant orientation (anisotropy) appears as peaks above this black curve. The color code refers to the developmental stages defined in the previous figures.

### Curvature

The virtual sections allowed us to study cell volumes and orientations across the spur (among sectors and between the lower and upper sides). A difference between both sides appears at stage 07, and a lower cell volume is observed in the upper side at stage 07 for all sectors. No particular trend is detected among developmental stages and among sectors during development (Supplementary Fig. S4).

## Discussion

Studies on spur morphogenesis in angiosperms remain scarce and have mainly concerned three unrelated genera, namely *Aquilegia* (Ranunculaceae), *Linaria* (Plantaginaceae) and *Centranthus* (Caprifoliaceae). These studies are focused on the epidermal cells, or are based on two-dimensional observations of this organ, often addressing the question of the origin of the interspecific diversity within a genus (*e.g.* Puzey *et al*., 2012; Galipot *et al*., 2021; Edwards *et al*., 2022). Comprehensive 3-dimensional information on all cells of the spur (not only the epidermal cells) is missing, hindering our full understanding of the mechanisms involved in complex petal forms. The present study aimed at filling this gap, using a new open source method of image processing developed for the purpose of this study. We observe that the development of the spur of *Staphisagria picta* proceeds in two phases. The first phase leads to the formation of an invagination by cell proliferation, whereas the second phase leads to spur elongation by anisotropic cell expansion in two different directions. The final shape, *i.e*. a thin and curved spur, results from the presence of cells oriented towards the spur cavity from the start of elongation and, cells that remain smaller and more spherical than others towards the distal part of the spur throughout development.

### Pattern of cellular processes during spur development

Our results on the development of the spur in *Staphisagria picta* suggest a pattern marked by an early phase of dominant cell proliferation that allows the formation of an invagination which is deeper than wide and that may already be defined as a spur. Subsequently, a second phase of directional (anisotropic) cell expansion dominates in most parts of the spur. These results are similar to those obtained in *Aquilegia* species that have spurred petals, suggesting convergence in cellular processes: a first phase of cell proliferation that stops early during development, followed by a longer phase of anisotropic cell elongation. However, it is difficult to go deeper in the comparison because the studies in *Aquilegia* focused on epidermal cells (Puzey *et al*., 2012) while we got data for the whole spur volume. In particular, our spatial analysis of spur elongation along three sectors shows that this elongation is not uniform within the spur. Cell expansion is concentrated in the proximal and median parts of the spur, which have the largest width. The cells are smaller and more spherical in the distal sector of the spur, corresponding to the tip. The second phase of development is marked by cell expansion, present everywhere in the organ, except in the most distal part. Moreover, cells have specific orientations throughout development. Cells oriented along the longitudinal axis of the spur are present at all stages, except stages 01 and 10, suggesting cell elongation. A population of cells oriented orthogonally to the longitudinal axis of the spur is also present, which may suggest thickening of the petal combined with enlargement of the opening and internal cavity of the spur in its upper part.

### Curvature

The detailed analysis of epidermal cells in *Aquilegia* petals bearing curved spurs (such as in *A. canadensis*) revealed that there is a differential growth between the distal and proximal surfaces of the spurs (relatively to the center of the flower), essentially due to differential cell division (Edwards *et al*., 2022). Our results do not support the hypothesis of a similar differential growth of the curved spur of *Staphisagria* when all cells are considered. However, they do reveal some complexity in the formation of the curvature since two cell populations with orthogonal orientations (along the longidutinal axis of the spur as well as orthogonal to this axis) are revealed at stages when curvature is achieved. This might be explained by differential tissue formation and growth between the epidermis and the parenchyma, suggesting different mechanisms in curvature formation in *Staphisagria* and *Aquilegia*.

### Vasculature

At stage 03, when the petal presents a flat blade and an emerging depression, three lines of elongated cells oriented from the proximal part to the distal part of the petal are observed. These elongated cells could be indicative of the vascular system. Petal vascularization was studied in three species of Delphinieae (*Aconitum lasiocarpum, Delphinium elatum, D. consolida*) (Novikoff and Jabbour, 2014), and in one species of Nigelleae (*Nigella damascena*) (Deroin *et al*., 2015), and the petals of all studied species have three vascular bundles. These cells are visible in the raw confocal microscopy images. In the three-dimensional reconstructions and in the volume distribution graphs, they are only visible in stage 03. This is due to the increase in the number of parenchyma cells that makes the cells of the vascular bundle undetectable. The observation of the confocal microscopy images shows that these lines of elongated cells become more numerous and branched as the spur grows, with the highest number of cells observed at stage 06. We hypothesize that this stage is characterized by high cell proliferation, and formation and branching of the vascular system. The vessels are tubular and made of lignified dead cells. This could explain why, at the subsequent stages, the walls could no longer be detected by the software with, as a consequence, a dramatic drop in the number of cells recognized.

### Diversity of spurs and short invaginations in Ranunculaceae and other eudicots

Convergence in spur development between *Aquilegia* and *Staphisagria* appears through the successive contribution of cell proliferation (mostly shape elaboration) and anisotropic cell elongation (deepening). However, more detailed comparative analyses have still to be conducted to determine the relative temporal contribution of the two phases to final shape in different species. In American *Aquilegia species*, spurs present a large diversity in length associated with the type of pollinators, and it has been shown that the duration of the phase of anisotropic elongation largely accounts for such diversity (Puzey *et al*., 2012). Actual spurs, *i.e.* deeper than wide invaginations, evolved only twice in Ranunculaceae but other forms of invaginations have been described in the family, in distantly related genera. This is the case in *Aconitum* (Delphinieae), *Nigella* (Nigelleae), *Urophysa* (Isopyreae), *Actaea* (Cimicifugeae), and *Coptis* (Coptideae) (reviewed in Delpeuch *et al*., 2022). These invaginations are variable in depth and width. Modeling petal shape based on a small set of morphogenetic parameters could account for this diversity (Cheng *et al*., 2023). The petals of *Nigella* species are spurless, but their blade is deeply invaginated and present two lips. At the cellular level, the development of the invagination of the petal of *Nigella damascena* is organized in two successive steps of cell proliferation and cell expansion, as observed in *Staphisagria* and *Aquilegia*. The first phase is responsible for most of the shape (Galipot *et al*., 2021). The two lips are formed by the differential expression domains of adaxial and abaxial genes during morphogenesis (Yao *et al*., 2019; Cheng *et al*., 2023). Adjusting the values of the morphogenetic parameters makes it possible to create the various petal shapes found in the genus (Cheng *et al*., 2023). How and if the morphogenesis of the invagination could relate to spur formation in the sister group Delphinieae remains to be explored in detail.

In the non-Ranunculaceae species in which spur development was studied, the relative contribution of each type of cellular processes varies. In *Centranthus ruber*, spur growth initially involves diffuse cell divisions, and then, from 30% of its final length until anthesis, cell elongation is mainly responsible for spur extension. Thus, it is a period of anisotropic cell elongation, with uniform elongation along the longitudinal axis of the spur, that primarily contributes to the length of the mature spur (Mack and Davis, 2015). In *Linaria* (Plantaginaceae), although both processes are present, cell proliferation was shown to be the primary mechanism involved in spur growth, and responsible for the difference in spur length among different species (Cullen *et al*., 2018).

Genetic origin of spurs has been investigated in a handful of species in eudicots and suggest that different genes are implicated. In *Aquilegia*, transcription factors of different families (TCP, ARF, C2H2 Zinc Finger) play a role in spur formation, some of which involved in auxin signaling (Yant *et al*., 2015; Ballerini *et al*., 2020; Zhang *et al*., 2020). In Delphinieae, the presence of a petal spur is closely linked to bilateral symmetry. A VIGS gene inactivation study in *Delphinium ajacis* has revealed interplay between a paralog of the petal identity gene *APETALA3-3* and a paralog of *CYCLOIDEA* (*CYL2b*) in the dorsal identity, including petal spur formation (Zhao *et al*., 2023). Transcription factor genes of the *KNOX* family have been suspected to direct spur formation in *Linaria* (Box *et al*., 2011).

### Methodological perspectives

Automated analysis now makes it possible to study organ morphogenesis in 3D, across all cell layers. This method of analysis allows the inclusion of large sample sizes and is therefore well suited to the study of developmental sequences. Producing confocal microscopy images transverse to the longitudinal axis of the spur would allow a more detailed study of the cellular mechanisms involved in the enlargement of the spur opening and cavity, and would also provide information on the formation of the curvature. It would also be interesting to separate the epidermal cells from the parenchymal cells to compare cellular processes between these tissues, and also to compare the results with those obtained from the study of the epidermis. The automation of the analyses allows now to carry out, in a more user-friendly way, 3D studies of organ morphogenesis on all cell layers in the framework of a developmental sequence, and including a larger number of samples.

## Conclusion

The way in which similar morphological structures are obtained through evolutionary convergence is a fascinating question. Here, we address this question by focusing on floral spurs. These structures are tightly associated with reproduction, by being invaginations in which nectar accumulates, providing a resource for potential pollinators. The analysis of the spur of *Staphiagria picta* revealed a process marked by an early phase of dominant cell proliferation, followed by a phase of anisotropic (directional) cell expansion, revealing that the convergence in form between the spurs of *Staphisagria* and *Aquilegia* is obtained by partially similar development processes. The method developped in the present study allowed precise 3D comparative studies at the cellular level. The recognition, identification, and extraction of information can be performed for each cell in a sample of organs of various sizes, typically belonging to a developmental sequence, using the same segmentation method. This allowed us to obtain a large amount of data and to perform a comprehensive 3-dimensional analysis of the morphogenesis of a complex structure. The same methodology is applicable to other complex organs and structures.

## Acknowledgements

We thank the seed bank of the French National Museum of Natural History (MNHN) for providing seeds of *Staphisagria picta*, and to the Jardin Botanique de Launay (Orsay, France) for the access to the greenhouse facility. We are grateful to IJPB-INRAE for providing access to the imaging facility, and to the Plateau Technique de Microscopie Électronique et de Microanalyses du MNHN (Géraldine Toutirais and Sylvain Pont).

## Author contributions

PD, SN and FJ designed the study. PD and KB performed the developmental analysis. PD and AP designed the pipeline for 3D image processing. PD wrote the first draft of the manuscript, SN, FJ and CD contributed to the writing. All authors contributed to the article and approved the submitted version.

## Data availability

The data underlying this article are available in the article and in its online supplementary material.

## Supplementary data

<u>Supplementary Figure S1:</u> Developmental benchmark for developmental stages of *Staphisagria picta* flowers. Figure adapted from (Delpeuch *et al.,* 2022). Blue, yellow and red colours refer to stamen, carpel, and petal development, respectively. The numbers in the purple disks correspond to the developmental stages targeted in the present study.

<u>Supplementary Figure S2:</u> Visualization of vascular tissue and parenchyma in images generated by confocal microscopy at stages 03 (top) and 09 (bottom). Yellow and blue arrows indicate respectively vascular tissue and parenchyma cells.

<u>Supplementary Figure S3:</u> Visualization of epidermal cells, cells on longitudinal and tangential sections, and vascular tissue generated by scanning electron microscopy at stage 07 and on mature petals. Blue and yellow arrows indicate respectively elongated cells and cells potentially oriented perpendicularly to the cutting plane.

<u>Supplementary Figure S4:</u> Study of the curvature of the spur. (A) Variations of cell volumes according to sectors and faces. The black spot in the boxplot is the average. (B) Variations of cell orientations. Plots represent the deviation angle of the directional axis of cells from the direction of the longitudinal axis of the petal for the spur. The small grid, on top left, indicates the position of the measurement. The color code refers to the developmental stages defined in the previous figures. The orange dot indicates the insertion point of the organ on the floral receptacle.

